# Revision of Paraglomeromycetes unveils a new order (Pervertustales), a new family (Innosporaceae), and a new genus (*Opikia*) in Glomeromycota

**DOI:** 10.64898/2026.09.12.751156

**Authors:** Piotr Niezgoda, Janusz Błaszkowski, Fritz Oehl, Ewald Sieverding, Thays Gabrielle Lins de Oliveira, Daniele Magna Azevedo de Assis, Bruno Tomio Goto, Mike Anderson Corazon-Guivin, Yongjun Liu, Gladstone Alves da Silva

**Author notes:** Piotr Niezgoda and Janusz Błaszkowski shared the first authorship of the paper due the equal contribution to the manuscript.

## Abstract

Currently, the fungal class Paraglomeromycetes (phylum Glomeromycota) consists of one order, Paraglomerales, with two families: Paraglomeraceae, comprising the genera *Paraglomus* with ten species and *Innospora* with one species, and the monogeneric and monospecific Pervetustaceae represented by *Pervetustus* with *Pe. simplex.* However, our morphological analyses of these species and an undescribed specimen morphologically resembling other Paraglomeromycetes species, as well as analyses of sequences of these fungi and environmental sequences, indicated that the current taxonomic composition of this class should be updated. We did so here by creating in Paraglomeromycetes Pervetustales ord. nov. with *Pervetustus* and *Opikia catenata* gen. nov. and sp. nov., as well as transferring *Innospora* with *In. majewskii* to Innosporaceae fam. nov. These novelties were introduced primarily based on strong genetic divergences of the analyzed taxa, as many morphological and histochemical features of their spores and mycorrhizal structures overlap. Only *Op. catenata* possesses the unique trait of occasionally producing spores in chains. This mode of sporulation has also not been recognized in any other Glomeromycota species. Unlike species of Pervetustales and Innosporaceae, which produce single-walled spores, the subcellular spore structure of *Paraglomus* species contains one or two walls.

## Introduction

The class Paraglomeromycetes was derived by Oehl et al. (2011) from Glomeromycetes sensu Cavalier-Smith (1998), which at that time was the only class encompassing all known taxa of arbuscular mycorrhizal fungi (AMF), classified in the phylum Zygomycota. Before the creation of Paraglomeromycetes, AMF were first transferred from the family Endogonaceae to the newly created order Glomerales with the type family and genus Glomeraceae and *Glomus*, respectively (Morton and Benny 1990). Subsequently, Morton and Redecker (2001) created in Glomerales the monogeneric family Paraglomeraceae with *Paraglomus*, represented by *Pa. brasilianum* and *Pa. occultum*. Both species were originally described as *Glomus brasilianum* and *Gl. occultum* (Walker 1982; Spain and Miranda 1996) because they produced glomoid spores formed similarly to other *Glomus* species. The main reasons for transferring these two *Glomus* species to *Paraglomus* were the strong divergences of their nuc rDNA sequences and histochemical properties of mycorrhizal structures from those of other glomoid spore-producing species. Schüßler et al. (2001) transferred AMF to the newly created phylum Glomeromycota and distributed these fungi into four orders, including Paraglomerales with Paraglomeraceae and *Paraglomus*.

In addition to the features mentioned above, Oehl et al. (2011) and Corazon-Guivin et al. (2020) concluded that *Paraglomus* species are also distinguished by the production of two-walled spores that usually germinate through the spore wall rather than through the spore subtending hyphal lumen, as glomoid spores of other taxa generally do. However, Morton and Redecker (2001) observed germ tubes of *Pa. brasilianum* also emerged through the subtending hyphal lumen.

Currently, apart from Paraglomeraceae with *Paraglomus*, Paraglomerales also includes the monogeneric and monospecific family Pervetustaceae with *Pervetustus* and *Pe. simplex* and the monospecific genus *Innospora* with *In. majewskii* (Paraglomeraceae), which were introduced into this order by Błaszkowski et al. (2017). *Innospora majewskii* was originally described as *Pa. majewskii* (Błaszkowski et al. 2012). The number of Paraglomerales species described so far is twelve, all of which, except *Pa. albidum*, *Pa. lacteum*, and *Pa. turpe*, have a known phylogeny.

Literature data and our own observations indicate that delimiting and classifying taxa of Paraglomeromycetes based on spore characteristics is difficult and often inconsistent with the phylogeny of the taxa. Such numerous discrepancies occur, e.g., between the distribution of sequenced Paraglomeromycetes species in the morphological key by Corazon-Guivin et al. (2020) and the distribution of these species in their phylogenetic tree.

In order to, at least partially, explain the reasons for the inconsistencies discussed above, we decided to verify the available data on morphological and histochemical characters of this fungal group, compare them with those of other members of Glomeromycota, and identify those characters that may have more clear diagnostic significance. Furthermore, we aimed to update the phylogeny of Paraglomeromycetes by including potentially undescribed members of this class in phylogenetic analyses and taking into account recent developments in the classification of the Glomeromycota.

## Materials and Methods

### Morphological analyses

Specimens of almost all AMF species attributed to Paraglomeromycetes were examined, including old specimens (established prior to 1990), most of which were mounted on microscopic slides in lactophenol. The mounting media used in studies of all other specimens were mainly mixtures of polyvinyl alcohol, lactic acid and glycerol (PVLG) or PVLG and Melzer’s reagent (Brundrett et al. 1994). In addition, intact and crushed spores mounted in a mixture of lactic acid and water (1:1, v/v), Melzer’s reagent, and water, as suggested by Spain (2003), were occasionally analyzed. In these analyses, we preferred to use spores that were freshly extracted from single or multiple spore-derived single-species cultures, although spores from trap cultures and field soil samples were also used. All original species descriptions and published species emendations were also considered and thoroughly studied, especially for those species whose specimens were unavailable to us.

The morphological and histochemical features taken into account in the verification and comparative analyses were: (i) the number of spore walls, (ii) the site of initiation of formation, flexibility, thickness, and composition of the innermost component of the subcellular spore structure in one-walled spores, (iii) the presence of an ornamented component of the subcellular spore structure, (vi) the cohesion of the subcomponents (laminae) of the major structural component of the subcellular spore structure, and (v) the staining of the components of the subcellular spore structure in Melzer’s reagent.

In addition, we studied spores and root fragments of a potentially new species, preliminary named Isolate 537. The spores were extracted from trap and single-species pot cultures, while the root fragments came from single-species cultures only to determine the properties of mycorrhizal structures of the isolate. The Isolate 537 spores were originally extracted from one of the 28 trap cultures inoculated with mixtures of root fragments and rhizosphere soils of *Ammophila arenaria* (L.) Link. The plant species colonized dunes of the Bay of Gdańsk located near Gdańsk-Stogi (54°37’N 18°72’E) in northern Poland. The root-rhizosphere soil samples were collected by P. Niezgoda on 5 June 2022. The trap and single-species cultures were established and grown, spores were extracted, and mycorrhizal structures were stained as described previously (Błaszkowski et al. 2006, 2009). The single-species cultures were established using ca. 10‒30 spores. According to Sayre et al. (2020), the climate type in the Gdańsk-Stogi region is cool temperate moist.

The terminology of subcellular spore structure follows that of Walker (1983) and Błaszkowski (2012). Nomenclature of fungi and the authors of fungal names are from the MycoBank website https://www.mycobank.org. Analyses of the spore walls, germination structures, and mycorrhizal structures were performed using compound microscopes at 100– 1000× magnification.

### DNA Extraction, PCR, Cloning, and DNA Sequencing

Genomic DNA of Isolate 537 was extracted from ca. 5‒10 spores. The method of processing the spores prior to PCR, conditions and primers used for PCR to obtain 45S (= 18S (partial)+ ITS+28S (partial)) nrDNA sequences of the isolates were as those described by Krüger et al. (2009) and Błaszkowski et al. (2021). Cloning and sequencing of PCR products were performed following protocols described by Błaszkowski et al. (2021). The sequences were deposited in GenBank.

### Phylogenetic analyses

To reconstruct the AMF phylogenies considered here, two alignments were created: (i) 45S with 45S sequences and (ii) 18S with almost complete 18S nrDNA sequences, including the new AMF 45S sequences obtained in this study and the published sequences, in NCBI, of all Paraglomeromycetes members (Suppl. Spreadsheet S1). *Archaeospora trappei* was included as outgroup in both alignments.

The alignments were aligned in MAFFT (Katoh et al. 2019) using default parameters. Prior to phylogenetic analyses, the model of nucleotide substitution was estimated with ModelTest-NG (Darriba et al. 2020). Maximum Likelihood - ML (1000 bootstrap) analyses were performed using RAxML-NG with Felsenstein Bootstrap Proportion (FBP) and Transfer Bootstrap Expectation (TBE), (Stamatakis 2014; Kozlov et al. 2019; Edler et al. 2020).

To determine the genetic similarity/relationship between the new genus and other taxa analyzed here, maximum identity (MI) values were determined using BLASTn (Johnson et al. 2008).

Clade and node supports were considered strong, moderate, and marginal when ML-TBE and ML-FBP support values were 0.95–1.00 and 81–100 %, 0.90–0.94 and 70–80 %, and < 0.90 and 70 %, respectively. The phylogenetic trees were visualized in FigTree ver. 1.4.4 (http://tree.bio.ed.ac.uk/software/figtree/) and edited in CorelDRAW X6 (Corel Corporation 2012).

## Results and discussion

### General data and phylogeny

The 45S alignment contained 65 sequences and had 2,150 sites (including gaps), of which 944 were constant, 219 variable, but not parsimony informative, and 987 were parsimony informative. The 18S alignment consisted of 51 sequences and had 1,808 sites, (including gaps), of which 1,437 were constant, 96 variable, but not parsimony informative, and 275 were parsimony informative.

The tree topologies generated by ML analyses (RAxML-NG with FBP and TBE) of the 45S and 18S alignments were similar. The analyzed species, Isolate 537, and the environmental sequences of the ingroup were accommodated in two major clades at the rank of order (Figs. 1 and 2). Each of these clades was strongly supported in both analyses (> 95 %). In the trees generated by Błaszkowski et al. (2017) and Tedersoo et al. (2024), these two major clades also obtained full support.

**Fig. 1.**
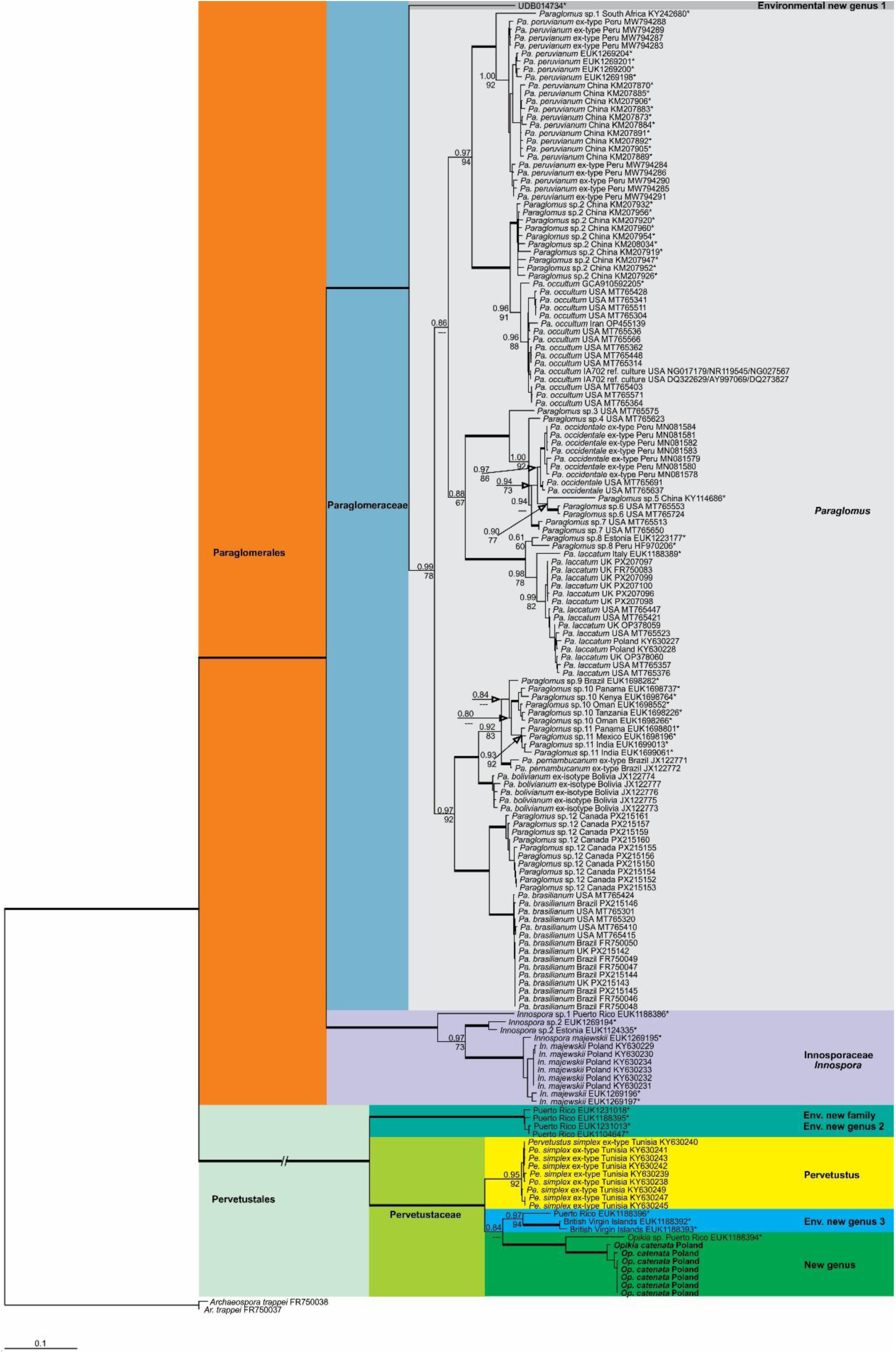
Phylogenetic tree obtained by analysis from partial 45S nrDNA sequences of Paraglomeromycetes. Sequences are labelled with their database accession numbers. Support values (from top) are from ML (maximum likelihood analysis) using RAxML-NG with TBE (Transfer Bootstrap Expectation) and FBP (Felsenstein Bootstrap Proportion), respectively. Only support values of at least 65 % are shown. Thick branches represent clades with more than 95 % of support in all analyses. Sequences obtained in this study are in boldface. The tree was rooted by sequences from *Archaeospora trappei*. The nucleotide substitution model used was GTR+I+G for both analyses.

**Fig. 2.**
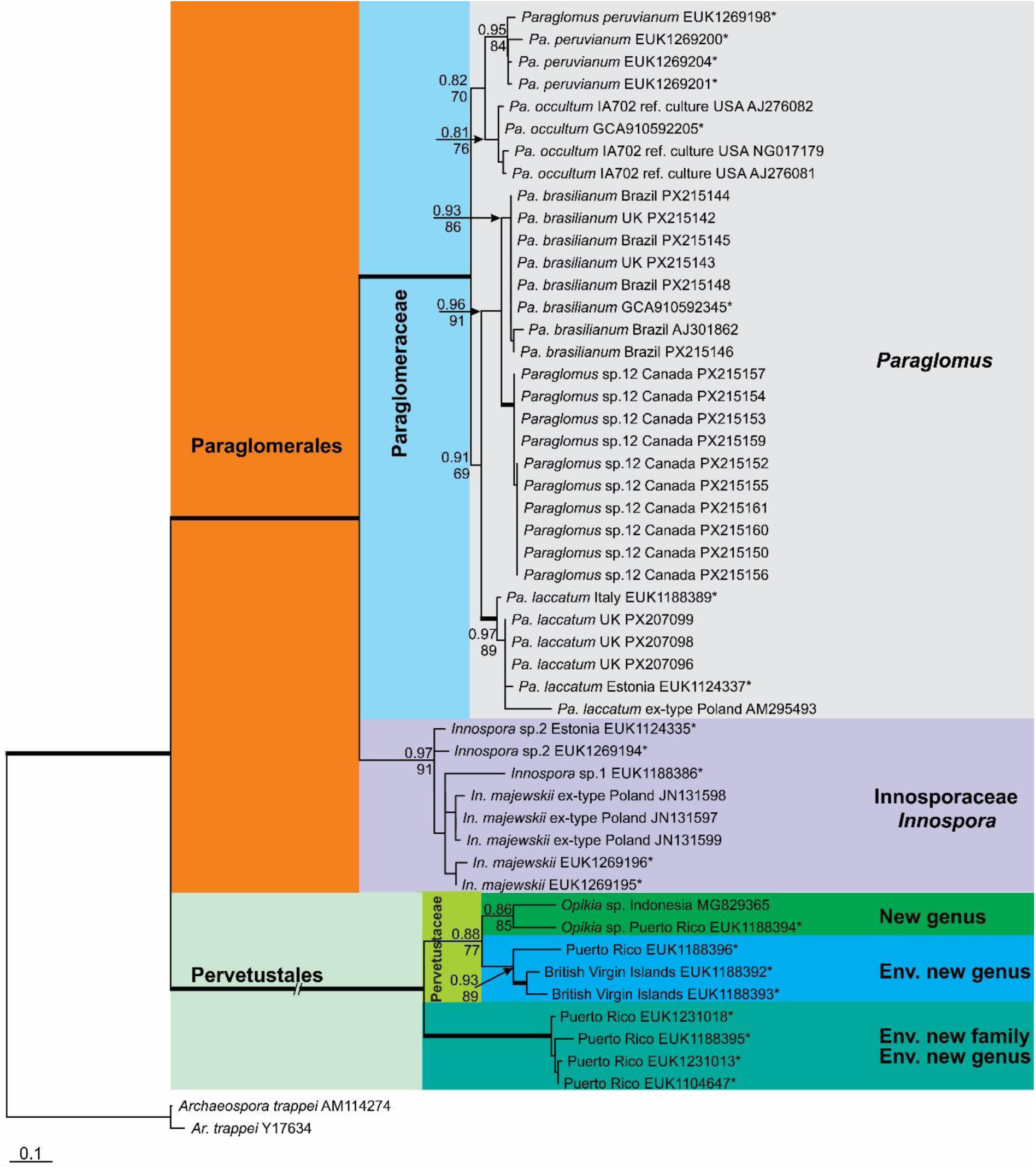
Phylogenetic tree obtained by analysis from almost complete 18S nrDNA sequences of Paraglomeromycetes. Sequences are labelled with their database accession numbers. Support values (from top) are from ML (maximum likelihood analysis) using RAxML-NG with TBE (Transfer Bootstrap Expectation) and FBP (Felsenstein Bootstrap Proportion), respectively. Only support values of at least 65 % are shown. Thick branches represent clades with more than 95 % of support in all analyses. The tree was rooted by sequences from *Archaeospora trappei*. The nucleotide substitution model used was GTR+I+G for both analyses.

In the 45S tree (Fig. 1), the first order clade clustered four environmental sequences in a clade at the rank of family, the family Pervetustaceae with (i) *Pervetustus* and *Pe. simplex*, (ii) Isolate 537, placed in a new generic clade, and (iii) three environmental sequences placed in another undescribed clade at the rank of genus. All these clades obtained high (≥ 0.95 and 92 %) or full support in both analyses.

The second clade at the rank of order contained two strongly supported (≥ 97 % in both analyses) clades at the rank of family, of which the first was inhabited only by *In. majewskii* and three environmental sequences representing two virtual taxa at species level (Fig. 1). The second family clade comprised one separated environmental sequence of a virtual taxon at the rank of genus and the genus *Paraglomus* with the species *Pa. bolivianum*, *Pa brasilianum*, *Pa. laccatum*, *Pa. occidentale*, *Pa. occultum*, *Pa. pernambucanum*, *Pa. peruvianum*, and 12 virtual taxa at species level.

The 18S tree generally reflected the ranges of taxon relationships shown in the 45S tree (Fig. 2), despite the absence of some components of this tree. Namely, this tree also consisted of two clades at the rank of order, each of which obtained high support values (≥ 95 %). *Pervetustus simplex* and Isolate 537 were not included in the 18S analyses because their 18S sequences are too short to provide reliable phylogenetic information. Therefore, we used much longer 18S sequences in our analyses, which were extracted from environmental 45S sequences clustering with 45S sequences representing Pervetustaceae. These analyses also placed Pervetustaceae in a basal clade at the rank of order and revealed the potential presence of two undescribed genera in Pervetustaceae, as well as a new family with a new genus in Pervetustales.

The second clade at the rank of order included two family clades, each of which obtained strong support (≥ 0.97 and 91 %) in both analyses (Fig. 2). The first clade, sister to a clade containing all *Paraglomus* species provided with 18S sequences, was inhabited by five *In. majewskii* sequences, including two coming from environmental analyses, and three environmental sequences that represented two potentially new species.

The second family clade consisted of the genus *Paraglomus*, represented by the species *Pa. brasilianum*, *Pa. laccatum*, *Pa. occultum*, and *Pa. peruvianum*. The 18S tree does not include *Pa. bolivianum*, *Pa. occidentale*, and *Pa. pernambucanum* because there are no available 18S sequences, including environmental sequences, for these species. *Paraglomus peruvianum* was represented by four environmental sequences with similarity suggesting conspecificity with this species. One virtual taxon, at species level, was represented by sequences near to *Pa. brasilianum*.

Some works for Glomerales and Ambisporales (Corazon-Guivin et al., 2019; Silva et al., 2023, 2025, 2026) demonstrated that differences among clades, ≥ 10 % in Maximum Identities (MI), considering the partial nrDNA gene, can indicate different genera. The MI, determined by BLASTn analyses with 45S sequences, showed that the most similar sequence to Isolate 537 sequences belongs to *Pe. simplex* (KY630244) with MI = 87.8 %. The sequences of the Isolate 537 differed from the *In. majewskii* KY630229 and *Pa. laccatum* MT765505 sequences, the latter representing *Paraglomus*, by at least 15.1 % and 15 %, respectively, indicating that Isolate 537 should represent a new genus.

### Morphological analyses

The verification and comparisons of the morphological and histochemical characters of the sequenced Paraglomeromycetes species revealed clear correlations between these characters and the phylogenetic position of these species shown in the tree illustrated in Fig. 1. Of the characters included in these analyses (see above, “Materials and Methods - Morphological analyses”), those of diagnostic importance are presented in the characteristics of the taxa at the ranks of genus to order (see below).

### Taxonomy

The phylogenetic and morphological analyses, as well as sequence comparisons described above, indicated that Paraglomeromycetes deserve a revision with a description of (i) a new order, based on Pervetustaceae, (ii) a new family, based on *Innospora majewskii*, and (iii) Isolate 537, which represents a new species belonging to an undescribed genus, which is most closely related to *Pervetustus*. The descriptions and data are given below.

### Paraglomeromycetes

Oehl, G.A. Silva, B.T. Goto & Sieverd., Mycotaxon 116: 374 (2011) MycoBank no.: MB 519687

Description: see Oehl et al. 2011 and below for the new subclass

Type subclass: *Paraglomeromycetidae* Oehl, Sieverd., G.A. Silva, Niezgoda, B.T. Goto & Błaszk.

### Paraglomeromycetidae

Oehl, Sieverd., G.A. Silva, Niezgoda, B.T. Goto & Błaszk., **subcl. nov.** MycoBank no.: MB 865203

Description: Spores formed terminally on hyphae; germination, as known so far, generally directly through spore wall; arbuscular mycorrhiza without or with only faint reaction when exposed to Trypan blue; vesicle formation unknown.

Etymology: Greek, *Para =* similar to, *glomus =* ball of yarn, referring to the morphological similarity with *Glomus* species.

Type order: *Paraglomerales* C. Walker & A. Schüssler

*Other order: Pervetustales* Błaszk., Oehl, Sieverd., B.T. Goto, Niezgoda & G.A. Silva

### Paraglomerales

C. Walker & A. Schüssler, Mycological Research 105 (12): 1418 (2001); emended here by Błaszk., Niezgoda, Sieverd., G.A. Silva, B.T. Goto & Oehl MycoBank no.: MB 90561

Emended description: Forming spores singly at tips of subtending hyphae. Spores colorless or lightly pigmented. Subcellular spore structure with three to six layers grouped in one wall, or in two walls: an outer and an inner wall. The inner wall has no physical contact with the outer wall and is formed *de novo* after the outer wall has fully differentiated. Outer spore wall layers with or without ornamentation. The main structural laminate spore wall layer with tightly adherent or easily separable laminae, not swelling in PVLG. Subtending hyphal wall composed of layers continuous with the layers of the outer spore wall or lacking the innermost spore wall layer in one-walled spores. Pore open or closed by (i) thickening layers of the subtending hyphal wall, (ii) the innermost layer of the outer spore wall, and (iii) a bridging septum continuous with the innermost laminae of the main structural laminate layer of the outer spore wall. Forming mycorrhiza with arbuscules, as well as intraradical and extraradical hyphae staining at most faintly or clearly in 0.1 % Trypan blue.

Type family: *Paraglomeraceae* J.B. Morton & D. Redecker

MycoBank no.: MB 82113

Other family: *Innosporaceae* G.A. Silva, Oehl, Sieverd., Niezgoda, B.T. Goto & Błaszk.

### Paraglomeraceae

J.B. Morton & D. Redecker, Mycologia 93 (1): 188 (2001); emended here by Błaszk., Niezgoda, Sieverd., G.A. Silva, B.T. Goto & Oehl

MycoBank no.: MB 82113

Emended description: Forming spores singly at tips of subtending hyphae. Spores colorless or lightly pigmented. Subcellular spore structure with three to six layers grouped in one or two walls: an outer wall and an inner wall. The inner wall has no physical contact with the outer wall and is formed *de novo* after the outer wall has fully differentiated. Outer spore wall layers with or without ornamentation. The main structural laminate spore wall layer with tightly adherent or easily separable laminae, not swelling in PVLG. Subtending hyphal wall composed of layers continuous with all layers of the spore wall forming the spore surface. Pore open or closed by (i) thickening layers of the subtending hyphal wall, (ii) the innermost layer of the outer spore wall, and (iii) a bridging septum continuous with the innermost laminae of the main structural laminate layer of the outer spore wall. Forming mycorrhiza with arbuscules, as well as intraradical and extraradical hyphae staining at most faintly in 0.1 % Trypan blue.

Type genus: *Paraglomus* J.B. Morton & D. Redecker

### Paraglomus

J.B. Morton & D. Redecker, Mycologia 93 (1): 188 (2001); emended here by Błaszk., Niezgoda, Sieverd., G.A. Silva, B.T. Goto & Oehl

MycoBank no.: MB 28458

Emended description: as for the family (here above)

Type species: *Paraglomus occultum* (C. Walker) J.B. Morton & D. Redecker, Mycologia 93 (1): 190 (2001)

MycoBank no.: MB 467740

Basionym: *Glomus occultum* C. Walker, Mycotaxon 15: 50 (1982)

MycoBank no.: MB 109779

### Innosporaceae

G.A. Silva, Oehl, Sieverd., Niezgoda, B.T. Goto & Błaszk., **fam. nov.** MycoBank no.: MB 865204

Description: Forming spores singly at tips of subtending hyphae. Spores colourless. Subcellular spore structure with one wall, the spore wall, consisting of three layers, of which the outermost layer 1, forming the spore surface, is short-lived, and the innermost layer 3 is thin (ca. 0.5 µm thick) and arises above or at the spore base at most. Subtending hyphal wall composed of two layers continuous with spore wall layers 1 and 2. Pore between the subtending hyphal lumen and the spore interior open. Forming mycorrhiza with arbuscules, as well as intraradical and extraradical hyphae staining clearly in 0.1 % Trypan blue.

Etymology: Latim, *inno* = float, *spora* = spore, referring to the spores of the type species floating in water.

Type genus: *Innospora* Błaszk., Kovács, Chwat & Kozłowska

### Innospora

Błaszk., Kovács, Chwat & Kozłowska, Nova Hedwigia 105 (3-4): 401 (2017) MycoBank no.: MB 820124

Description: as for the family (here above) and see Błaszkowski et al. (2017)

Type species: *Innospora majewskii* (Błaszk. & Kovács) Błaszk., Kovács, Chwat & Kozłowska, Nova Hedwigia 105 (3-4): 403 (2017)

### MB 820125

Basionym: *Paraglomus majewskii* Błaszk. & Kovács, Mycologia 104 (1): 149 (2012) MycoBank no.: MB 561205

### Pervetustales

Błaszk., Oehl, Sieverd., B.T. Goto, Niezgoda & G.A. Silva, **ord. nov.**

MycoBank no.: MB 865205

Description: Producing spores singly, in loose clusters at tips of subtending hyphae, and in short chains with two (-three) spores, the first of which is formed at the tip of the subtending hypha continuous with an extraradical mycorrhizal hypha, and the second (third) spore is formed at the tip of a short hypha, with a wall comprising layers continuous with the spore wall layers of the first (second) spore. Spores hyaline or clearly pigmented. Subcellular spore structure with one spore wall containing two to five layers. The main structural spore wall layer, 11 the laminate layer, always swells slightly in PVLG. Subtending hyphal wall composed of layers continuous with spore wall layers. Pore open or closed by one or two septa bridging the innermost subtending hyphal wall layer. Forming mycorrhiza with arbuscules, as well as intraradical and extraradical hyphae staining faintly to greyish violet in 0.1 % Trypan blue.

Etymology: Latim, *pervetustus* = very old, refering to the very old origin based on molecular phylogeny.

Type family: *Pervetustaceae* Błaszk., Chwat, Kozłowska, Symanczik & Al-Yahya’ei

### Pervetustaceae

Błaszk., Chwat, Kozłowska, Symanczik & Al-Yahya’ei, Nova Hedwigia 105 (3-4): 403 (2017)

MycoBank no.: MB 820126

Description: see Błaszkowski et al. (2017) and the description of Pervetustales (see above)

Type genus: *Pervetustus* Błaszk., Chwat, Kozłowska, Symanczik & Al-Yahya’ei

Other genus: *Opikia* Błaszk., Niezgoda, B.T. Goto, Sieverd., Oehl & G.A. Silva

### Pervetustus

Błaszk., Chwat, Kozłowska, Symanczik & Al-Yahya’ei, Nova Hedwigia 105 (3-4): 404 (2017)

MycoBank no.: MB 820127

Description: see Błaszkowski et al. (2017)

Type species: *Pervetustus simplex* Błaszk., Chwat, Kozłowska, Crossay, Symanczik & Al-Yahya’ei, Nova Hedwigia 105 (3-4): 404 (2017) MycoBank no.: MB 820128

### Opikia

Błaszk., Niezgoda, B.T. Goto, Sieverd., Oehl & G.A. Silva, **gen. nov.** Figs. 1, 2, 3A‒H, 4A‒D

**Fig. 3.**
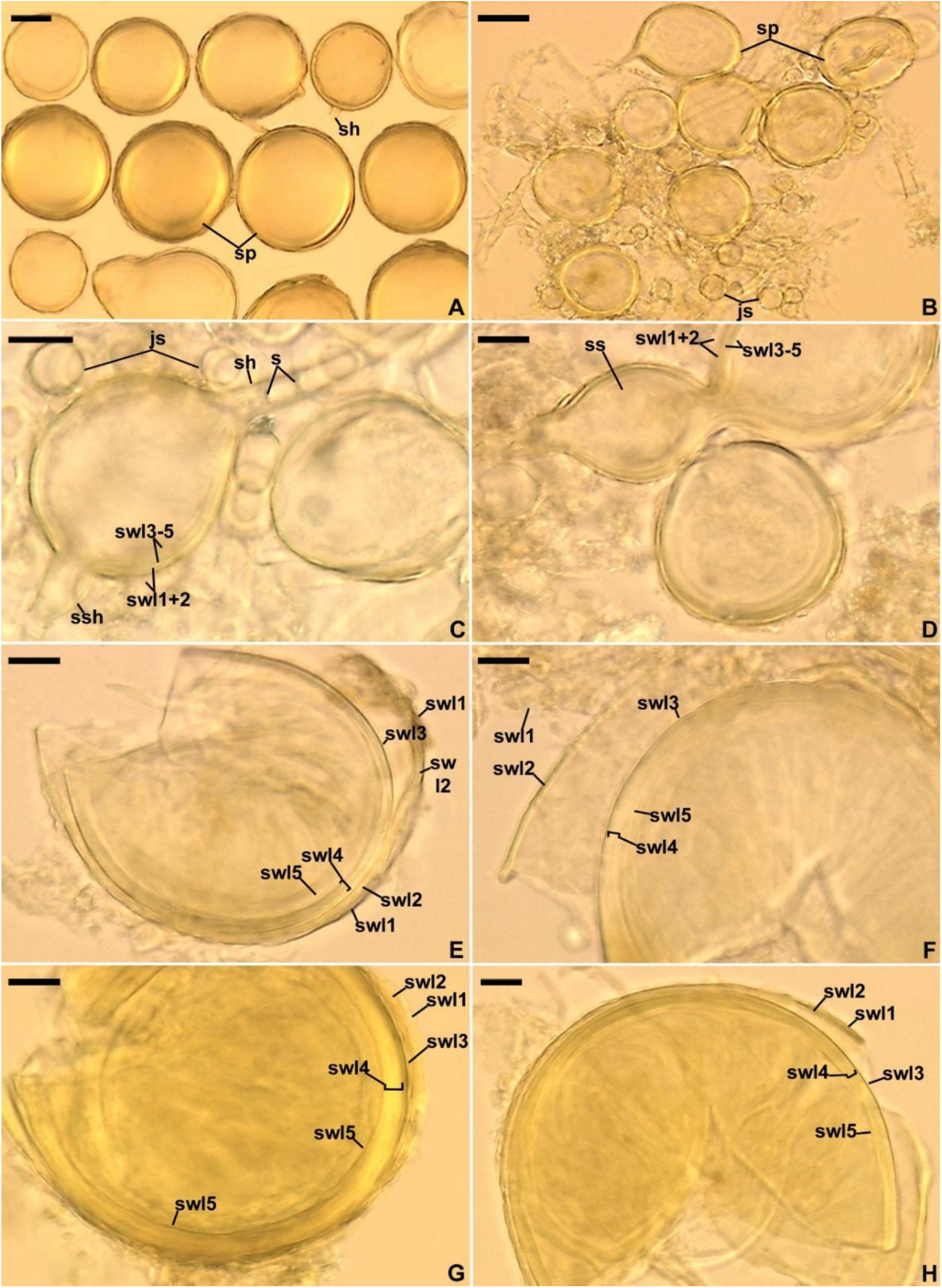
Opikia catenata. **A.** Spore (sp) produced singly; subtending hypha is indicated. **B.** Spores (sp) in a loose cluster; juvenile spores (jsp) are indicated. **C.** Spore with a five-layered spore wall (swl1‒5), subtending hypha (sh) with cross septa (s), and a secondary spore (ssp) beginning to develop from the parent spore. **D.** Spore wall layers (swl) 1‒5 of the parent spore continuous with spore wall layers of the secondary spore. **E‒H.** Spore wall layers (swl) 1‒5. **A.** Glomerocarps in lactic acid. **B‒F.** Spores in PVLG. **G, H.** Spores in PVLG+Melzer’s reagent. **A‒H.** Differential interference microscopy. Scale bars: **A** = 25 μm, **B** = 20 μm, **C‒H** = 10 μm.

MycoBank no.: MB 865206

Etymology: *Opikia*, in honor of Professor Maarja Öpik, University of Tartu, Estonia, for her enormous contribution to the knowledge of the ecology and distribution of AMF.

Type species: *Op. catenata* Błaszk., Niezgoda, B.T. Goto, Sieverd., Oehl & G.A. Silva

Diagnosis: Differs from (A) all other genera of Glomeromycota in the ability to produce spores in chains and (B) the monospecific *Pervetustus* with *Pe. simplex* in (i) the ability to form spores in loose clusters, (ii) size of spores, (iii) the subcellular spore structure and the phenotypic features of spore wall layer 1, (iv) morphometric characters of the spore subtending hypha, and (v) nucleotide composition of sequences of the 45S nuc rDNA region (see “Notes” for details).

Genus description: As that of *Opikia catenata* (see below).

### Opikia catenata

Błaszk., Niezgoda, B.T. Goto, Sieverd., Oehl & G.A. Silva, **sp. nov.** Figs. 3A‒ H, 4A‒D

MycoBank no.: MB 865381

Etymology: *catenata*, Latin, referring to the production of spores in chains by the new species.

Typification: POLAND. Pomerania Voivodeship, glomerospores (= spores) from a single-species culture extracted from a trap culture inoculated with the rhizosphere soil and root fragments of *Am. arenaria* from dunes of the Bay of Gdańsk located near Gdańsk-Stogi (54°37’N 18°72’E) in norther Poland, 5 June 2022, P. Niezgoda (**holotype**: slide with spores Z+ZT Myc 0094525).

Diagnosis: Differs from *Pe. simplex*, the closest phylogenetic relative (Fig. 1) in (i) the ability to form spores in loose clusters and chains, (ii) size of spores, (iii) the subcellular spore structure and the phenotypic features of spore wall layer 1, (iv) morphometric characters of the spore subtending hypha, and (v) nucleotide composition of sequences of the 45S nuc rDNA region (see “*Notes*” for details).

Description: Forming glomoid spores (i) singly, at tips of subtending hyphae continuous with extraradical mycorrhizal hyphae, (ii) in loose clusters, at tips of subtending hyphae branched from a parent hypha continuous with an extraradical mycorrhizal hypha, and (iii) in short chains with two (‒three) spores, the first of which is formed at the tip of the subtending hypha continuous with an extraradical mycorrhizal hypha, and the second (third) spore is formed at the tip of a short hypha, with a wall comprising layers continuous with the spore wall layers of the first (second) spore, grown opposite the base of the first spore (Figs. 3A‒H, 4A‒ C). *Spores* yellowish white (4A2) to brownish yellow (5C7); globose to subglobose; (36‒)58(‒ 88) µm diam; occasionally ovoid to oblong; 54‒60 × 60‒85 µm; with one subtending hypha (Figs. 3A‒H, 4A‒C). *Subcellular structure of spores* composed of one spore wall with one yellowish white (4A2) to brownish yellow (5C7), semi-permanent layer (layer 1) and four hyaline, permanent layers (layers 2‒5) (Figs. 3C‒H, 4A‒C). Layer 1, forming the spore surface, yellowish white (4A2) to brownish yellow (5C7), semi-permanent, friable, flexible to semi-flexible, (0.7‒)1.8(‒2.5) µm thick, easily separating from layer 2, slowly deteriorating with age, usually present as a more or less decomposed structure even in older spores, rarely completely sloughed off (Figs. 3C‒H, 4A‒C). Layer 2 hyaline, permanent, uniform (without visible sublayers), semi-flexible, (1.0‒)1.7(‒2.0) µm thick, easily separating from the upper surface of layer 3 (Figs. 3C‒H, 4A‒C). Layer 3 uniform, semi-flexible, (0.6‒)0.9(‒1.4) µm thick, tightly adherent to the upper surface of layer 4, not separating from this layer in even vigorously crushed spores (Fig. 3E‒H). Layer 4 laminate, semi-flexible, (2.0‒)3.1(‒4.4) µm thick, consisting of very thin, <0.5 µm, sublayers loosely connected to each other; the innermost sublayer(s) often clearly detach from the other sublayers in vigorously crushed spores, giving the false impression of the presence of a separate layer (Figs. 3E‒H, 4A, B); layer 4 always slightly swelling in PVLG. Layer 5 flexible, 0.6‒0.8 µm thick, often difficult to see because of its similarity to the sublayers of the laminate layer 4 and when not separated from the inner surface of that layer (Fig. 3E‒H). In Melzer’s reagent, layer 2 remains hyaline or stains pale yellow (3A3) and layer 4 turns pale yellow (4A3) to light yellow (4A4) (Figs. 3G, H, 4A). The reactivity of layer 5 in this reagent remains unknown because all attempts to extract this layer from the spore to be directly visible (not under the other layers) failed. *Subtending hypha* yellowish white (4A2) to brownish yellow (5C7); straight or recurved, cylindrical to funnel-shaped; (4.9‒)6.5(‒7.6) µm wide at the spore base (Figs. 3A, C, 4A‒C). *Wall of subtending hypha* yellowish white (4A2) to brownish yellow (5C7), rarely hyaline; (1.8‒)2.5(‒3.6) µm thick at the spore base; composed of five layers continuous with spore wall layers 1‒5 (Fig. 4A‒C). *Pore* (1.2‒)1.4(‒2.2) µm diam, open (Fig. 4C) or occluded by one to two, straight or slightly curved, septa bridging the inner surfaces of the innermost layer of the subtending hyphal wall; septa, 0.6‒0.8 µm thick, formed at or below the spore base (Figs. 3C, 4B). *Germination* unknown.

**Fig. 4.**
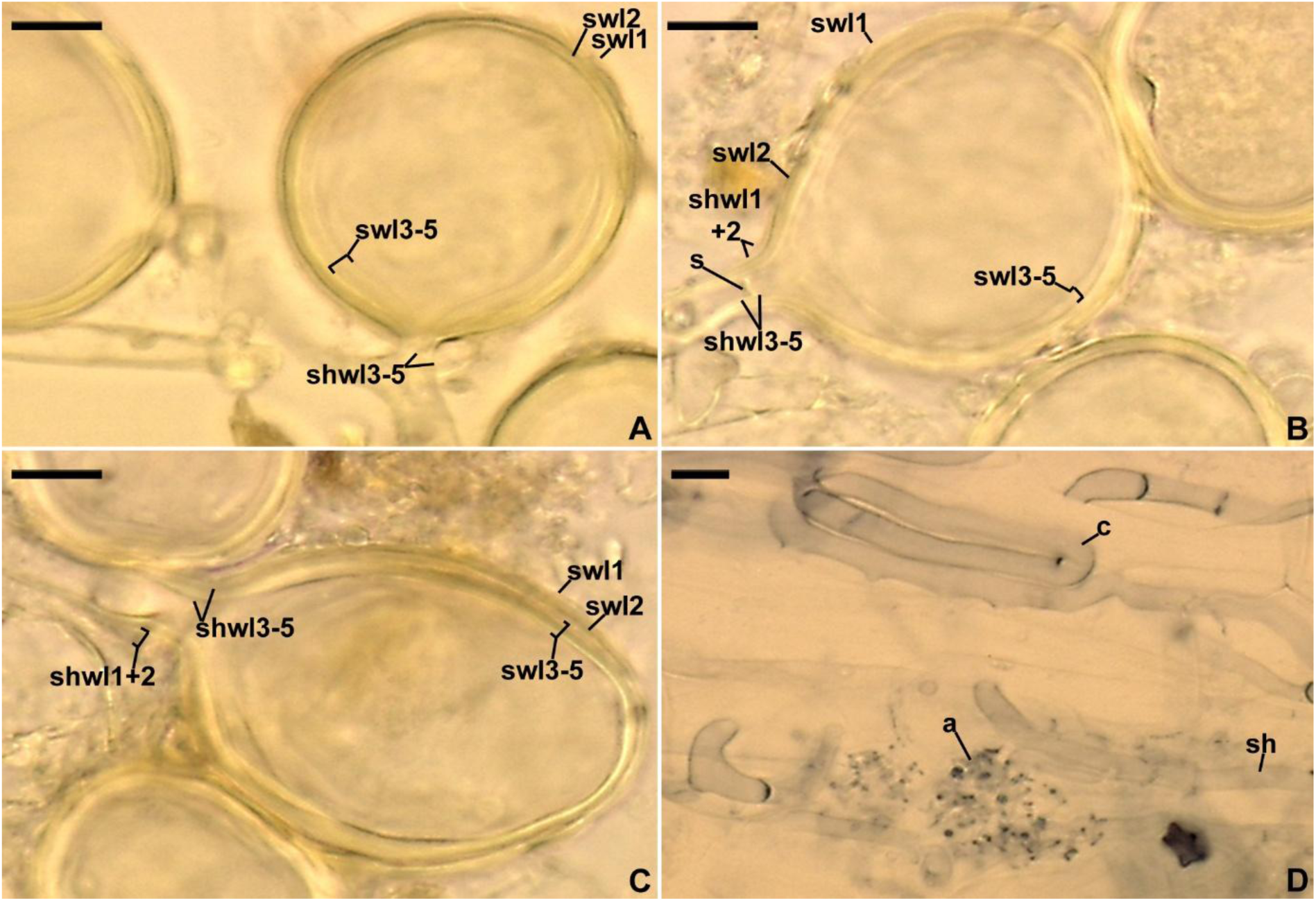
Opikia catenata. **A‒C.** Spore wall layers (swl) 1‒5 continuous with subtending hyphal wall layers (shwl) 1‒5. **D.** Arbuscule (a), coil (c), and straight hypha (sh) in a mycorrhizal root fragment stained in 0.1 % Trypan blue. **A.** Spores in PVLG+Melzer’s reagent. **B‒D.** Spores and mycorrhizal structures in PVLG. **D. A‒C.** Differential interference microscopy. Scale bars: **A‒D** = 10 μm.

Ecology and distribution: In the field, *Op. catenata* probably lived in mycorrhizal symbiosis with *Am. arenaria* that colonized dunes of the Bay of Gdańsk located near Gdańsk-Stogi (54°37’N 18°72’E) in northern Poland. However, no molecular analysis was performed to confirm this supposition. In single-species cultures with *Pl. lanceolata* as the host plant, *Op. catenata* formed mycorrhiza with arbuscules, as well as intraradical and extraradical hyphae staining faintly, violet white (16A2) to lilac grey (16B2), or not staining in 0.1 % Trypan blue. GenBank and EUKARYONE searches revealed many sequences with > 90 % identity to the 45S *Op. catenata* sequences. However, ML analyses showed that only 10 of these sequences with > 96 % identity clustered in a clade with the *Op. catenata* sequences (data not shown). The sequences were obtained from soils collected in South Africa, Estonia, Iran, Mexico, Russia, and the USA (for details see suppl. Table S1). Thus, *Op. catenata* probably has a wide distribution in the world.

Notes: The feature that makes *Op. catenata* a unique AM fungus at the species level and to all higher-ranking taxa of Glomeromycota is the production of spores in chains (Fig. 3C, D). In addition, the formation of spores in loose clusters by *Op. catenata* (Fig. 3B) is a trait not known in any other Paraglomeromycetes.

Morphologically, *Op. catenata* and *Pe. simplex*, the closest phylogenetic relatives (Fig. 1), differ primarily in the composition of the spore wall and phenotypic features of its components (layers). The spore wall of *Op. catenata* comprises five layers (Figs. 3C‒H, 4A‒C), while that of *Pe. simplex* consists of two layers (Błaszkowski et al. 2017). In *Op. catenata*, spore wall layer 1 is pigmented, degrades slowly with age, and is usually present, at least partially, in even older spores (Figs. 3C‒H, 4A‒C). In *Pe. simplex*, spore wall layer 1 is colorless, short-lived, and usually strongly deteriorated or completely sloughed off in newly mature spores. Importantly, the *Pe. simplex* spore wall layer 1 lacks the distinguishing feature of the *Op. catenata* spore wall layer 1, i.e., friability. In this respect, spore wall layer 1 of *Op. catenata* resembles spore wall layer 1 of *Ambispora leptoticha* (Archaeosporales) (Bills and Morton 2015), which, however, produces acaulosporoid spores, i.e., laterally on the neck of a sporiferous saccule like *Acaulospora* species. Finally, *Op. catenata* spores can be distinctly, up to 2.1-fold, larger when globose, and at the spore base their subtending hypha and the subtending hyphal wall are 1.3‒1.6-fold wider and 1.8‒2.0-fold thicker, respectively.

Genetically, *Op. catenata* and *Pe. simplex* also differ significantly. The 45S sequence divergences between these species reached 12.2‒14.2 %, clearly exceeding the 10 % threshold that separates most Glomeromycota genera (Corazon-Guivin et al., 2019; Silva et al., 2023; Silva et al., 2025; 2026).

Of the non-sequenced members of Paraglomerales, in morphology *Op. catenata* most closely resembles *Pa. turpe*. The main feature sharing these two species is their complex, multilayered, subcellular spore structure. While *Op. catenata* spores have five layers, clearly constituting one structure, the spore wall (Figs. 3C‒H, 4A‒C), Oehl et al. (2016) arranged spore layers of *Pa. turpe* in two walls, a three-layered outer wall and a two- to three-layered inner wall. In addition, spore wall layer 2 of *Op. catenata* is persistent (Figs. 3C‒H, 4A‒C), while all three layers of the outer wall of *Pa. turpe* rapidly deteriorate with age and are rarely present due to complete decomposition. Four other characters also clearly separate *Op. catenata* from *Pa. turpe*. Young and mature spores of *Op. catenata* are yellowish white (4A2) to brownish yellow (5C7) (Figs. 3A‒F, 4B, C), while *Pa. turpe* spores remain hyaline regardless of age. *Opikia catenata* spores can be up to 1.8-fold larger when globose to subglobose, and at the spore base the subtending hypha of the new species can be up to 1.5-fold wider and has a wall 1.6-fold thicker.

Taking into account the comparisons and information presented above, especially those showing taxonomically significant differences between *Op. catenata* and *Pa. turpe* in subcellular spore structure and the lack of available genetic sequences of *Pa. turpe*, we left this species in *Paraglomus*.

### Final notes

The results of the studies described here and those by Tedersoo et al. (2024), based on analyses of sequences of described species and environmental sequences, demonstrated that Paraglomeromycetes is a very species-poor class of Glomeromycota. Taking into account the *Op. catenata* sp. nov. described here, this class comprises 13 species, i.e., only ca. 3.4 % of all described species of the phylum. This fact is surprising because Paraglomeromycetes is considered the oldest group of AMF (Malar et al. 2022). Therefore, this group has had the longest time to develop diversity. So, why is the diversity of Paraglomeromycetes so low?

Three main causes probably led to this impoverishment. (1). The relatively small genome size and content of members of Paraglomeromycetes, as evidenced for *Pa. occultum* (Malar et al. 2022). This may have contributed to the low colonization potential of these fungi, resulting in a reduction in the number of successfully colonized host plants and the amount of nutrients utilized from plant resources. Consequently, this reduction impaired the reproductive performance of these fungi and the condition of their host plants, the latter by reducing the number of benefits acquired from the fungi (Säle et al. 2021). In addition, it weakened the ability of the fungal partners to compete with co-occurring soil organisms and adapt to harmful environmental influences. All these effects ultimately suppressed the development of genetic diversity in this fungal group and selected genotypes that tolerated these disadvantages. One such genotype is likely *In. majewskii*. Analyses of at least 10,000 trap cultures showed that it is one of the most common and frequently occurring Glomeromycota species in cultivated and uncultivated sites, including particularly onerous coastal dunes, in various regions of the world (Błaszkowski, pers. observ.). Furthermore, this resilience to nuisances is also reflected in the fact that *In. majewskii* is easily grown in single-species cultures, where it produces enormous numbers of spores. In contrast, the same analyses did not reveal any other Paraglomeromycetes species, with the exception of the generally rarely occurring *Pe. simplex* and *Op. catenata*. (2). The lack of recognition of much of the diversity of this fungal group due to at least three reasons. The first one is the low and often zero efficiency of primers amplifying DNA fragments with high species resolution for these fungi. Our numerous attempts to obtain *rpb1* and glomalin sequences of *Op. catenata* and other members of Paraglomeromycetes failed. The second reason is the exceptional difficulty of finding Paraglomeromycetes spores. The aforementioned *In. majewskii* was almost never observed in field-collected soil samples, despite sporulating abundantly in trap cultures inoculated with these samples. The invisibility or absence of spores of members of Paraglomeromycetes, most of which produce spores morphologically similar to those of *In. majewskii*, in field samples is certainly due to the difficulty in detecting them (they are very small and usually colorless) and short-lived nature. The latter is due to the presence of delicate components in the spore structure, easily degraded by soil hyperparasites. The third reason is the very small group of mycologists engaged in collecting, growing, and characterizing the morphology and phylogeny of Glomeromycota (Goto et al. 2024).

The high similarity of the topology of the Paraglomeromycetes trees generated by us and Tedersoo et al. (2024), and the high clade supports in these trees, including those for *In. majewskii* and *Pe. simplex* plus *Op. catenata*, prove that our description of the new taxa in this class, i.e., Innosporaceae, Pervetustales, and *Opikia*, is well justified.

Issues that remain unclear include the number of spore walls in *Paraglomus* species and the mode of spore germination of all members of Paraglomeromycetes. Resolving these uncertainties will require challenging ontogenetic analyses and studies aimed at determining the trajectory of germ tubes in at least several species of all higher-ranking taxa of this fungal group. Difficulties in these studies will stem from (i) the small size of spores, which have relatively few and difficult-to-observe components of the subcellular spore structure, (ii) the low success rate of growing these fungi in single-species cultures, and (iii) the lack of availability of living specimens of most Paraglomeromycetes members.

## Acknowledgments

Daniele Magna Azevedo de Assis and Thays Gabrielle Lins de Oliveira thanks the Fundação de Amparo à Ciência e Tecnologia do Estado de Pernambuco (FACEPE) for providing fellowship. Gladstone A. Silva and Bruno T. Goto has a fellowship from the Conselho Nacional de Desenvolvimento Científico e Tecnológico (CNPq) (Proc. 314045/2026-0; 302513/2026-4). Piotr Niezgoda and Yongjun Liu were supported by the Joint Research Project between Polish National Centre of Science (grant no. 2025/56/Q/NZ9/00414) and National Science Foundation of China (grant no. W2512021). This work was supported by INCT - Brazilian Fungi Project funded by Conselho Nacional de Desenvolvimento Científico e Tecnológico (CNPq) (proc. 409181/2024-2).

