## Supplementary material for "Revision of Paraglomeromycetes unveils a new order (Pervertustales), a new family (Innosporaceae), and a new genus (*Opikia*) in Glomeromycota": Spreadsheet S1

**Spreadsheet S1**. Accession numbers for the sequences used in the phylogenetic analyses of this study.

| Species name in NCBI | ID | nrDNA region | Country | Accession numbers (nrDNA) |
|---|---|---|---|---|
| *Innospora majewskii* | Ex-type | SSU | Poland | JN131597-JN131599 |
| *Innospora majewskii* | 1 | SSU+ITS+LSU | Poland | KY630229 |
| *Innospora majewskii* | 10 | SSU+ITS+LSU | Poland | KY630230 |
| *Innospora majewskii* | 6 | SSU+ITS+LSU | Poland | KY630231 |
| *Innospora majewskii* | 7 | SSU+ITS+LSU | Poland | KY630232 |
| *Innospora majewskii* | 8 | SSU+ITS+LSU | Poland | KY630233 |
| *Innospora majewskii* | 9 | SSU+ITS+LSU | Poland | KY630234 |
| *Innospora majewskii* env. sample | Not available | SSU+ITS+LSU | Not available | EUK1269195 |
| *Innospora majewskii* env. sample | Not available | SSU+ITS+LSU | Not available | EUK1269196 |
| *Innospora majewskii* env. sample | Not available | SSU+ITS+LSU | Not available | EUK1269197 |
| *Innospora* sp.1 env. sample | S050 | SSU+ITS+LSU | Puerto Rico | EUK1188386 |
| *Innospora* sp.2 env. sample | G5803 | SSU+ITS+LSU | Estonia | EUK1124335 |
| *Innospora* sp.2 env. sample | Not available | SSU+ITS+LSU | Not available | EUK1269194 |
| *Opikia catenata* | Ex-type | SSU+ITS+LSU | Poland | |
| *Opikia* sp. env. sample | Singleton | SSU | Indonesia | MG829365 |
| *Opikia* sp. env. sample | S045 | SSU+ITS+LSU | Puerto Rico | EUK1188394 |
| *Paraglomus* sp.6 | ASV430 | SSU+ITS+LSU | USA | MT765724 |
| *Paraglomus bolivianum* | ex-isotype | LSU | Bolivia | JX122773-JX122777 |
| *Paraglomus brasilianum* | Att260-4 | SSU | Brazil | AJ301862 |
| *Paraglomus brasilianum* | Att260-8 | SSU+ITS+LSU | Brazil | FR750046-FR750050 |
| *Paraglomus brasilianum* | ASV_7 | SSU+ITS+LSU | USA | MT765301 |
| *Paraglomus brasilianum* | ASV_26 | SSU+ITS+LSU | USA | MT765320 |
| *Paraglomus brasilianum* | ASV_116 | SSU+ITS+LSU | USA | MT765410 |
| *Paraglomus brasilianum* | ASV_121 | SSU+ITS+LSU | USA | MT765415 |
| *Paraglomus brasilianum* | ASV_130 | SSU+ITS+LSU | USA | MT765424 |
| *Paraglomus brasilianum* | W960_3_902 | SSU+ITS+LSU | UK | PX215142, PX215143 |
| *Paraglomus brasilianum* | W260_5_933C | SSU+ITS+LSU | Brazil | PX215144, PX215146, PX215148 |
| *Paraglomus brasilianum* | BR105_2435A | SSU+ITS+LSU | Brazil | PX215145 |
| *Paraglomus brasilianum* env. sample | Not available | SSU | Not available | GCA910592345 |
| *Paraglomus laccatum* | Ex-type | SSU | Poland | AM295493 |

| | | | | |
|---|---|---|---|---|
| *Paraglomus laccatum* | P4_bc1083_SPU_5_13 | SSU+ITS+LSU | UK | PX207097 |
| *Paraglomus laccatum* | P4_bc1083_SPU_5_13 | SSU+ITS+LSU | UK | PX207099 |
| *Paraglomus laccatum* | P4_bc1083_SPU_5_13 | SSU+ITS+LSU | UK | PX207100 |
| *Paraglomus laccatum* | P4_bc1083_SPU_5_13 | SSU+ITS+LSU | UK | PX207096 |
| *Paraglomus laccatum* | P4_bc1083_SPU_5_13 | SSU+ITS+LSU | UK | PX207098 |
| *Paraglomus laccatum* | W5141 | SSU+ITS+LSU | UK | FR750083 |
| *Paraglomus laccatum* | ASV_63 | SSU+ITS+LSU | USA | MT765357 |
| *Paraglomus laccatum* | ASV_82 | SSU+ITS+LSU | USA | MT765376 |
| *Paraglomus laccatum* | ASV_127 | SSU+ITS+LSU | USA | MT765421 |
| *Paraglomus laccatum* | ASV_153 | SSU+ITS+LSU | USA | MT765447 |
| *Paraglomus laccatum* | ASV_229 | SSU+ITS+LSU | USA | MT765523 |
| *Paraglomus laccatum* | W6534 | SSU+ITS+LSU | UK | OP378059, OP378060 |
| *Paraglomus laccatum* | 25 | SSU+ITS+LSU | Poland | KY630227 |
| *Paraglomus laccatum* | 26 | SSU+ITS+LSU | Poland | KY630228 |
| *Paraglomus laccatum* env. sample | S811 | SSU+ITS+LSU | Italy | EUK1188389 |
| *Paraglomus laccatum* env. sample | G5901 | SSU+ITS+LSU | Estonia | EUK1124337 |
| *Paraglomus occidentale* | Ex-type | SSU+ITS+LSU | Peru | MN081578-MN081584 |
| *Paraglomus occidentale* | ASV343 | SSU+ITS+LSU | USA | MT765637 |
| *Paraglomus occidentale* | ASV397 | SSU+ITS+LSU | USA | MT765691 |
| *Paraglomus occultum* | ASV10 | SSU+ITS+LSU | USA | MT765304 |
| *Paraglomus occultum* | ASV20 | SSU+ITS+LSU | USA | MT765314 |
| *Paraglomus occultum* | ASV47 | SSU+ITS+LSU | USA | MT765341 |
| *Paraglomus occultum* | ASV68 | SSU+ITS+LSU | USA | MT765362 |
| *Paraglomus occultum* | ASV70 | SSU+ITS+LSU | USA | MT765364 |
| *Paraglomus occultum* | ASV106 | SSU+ITS+LSU | USA | MT765403 |
| *Paraglomus occultum* | ASV134 | SSU+ITS+LSU | USA | MT765428 |
| *Paraglomus occultum* | ASV154 | SSU+ITS+LSU | USA | MT765448 |
| *Paraglomus occultum* | ASV217 | SSU+ITS+LSU | USA | MT765511 |
| *Paraglomus occultum* | ASV242 | SSU+ITS+LSU | USA | MT765536 |
| *Paraglomus occultum* | ASV272 | SSU+ITS+LSU | USA | MT765566 |
| *Paraglomus occultum* | ASV277 | SSU+ITS+LSU | USA | MT765571 |
| *Paraglomus occultum* | VRU62 | SSU+ITS+LSU | Iran | OP455139 |
| *Paraglomus occultum* | IA702 ref. culture | SSU | USA | AJ276081, AJ276082, DQ322629, NG017179 |

| | | | | |
|---|---|---|---|---|
| *Paraglomus occultum* | IA702 ref. culture | ITS | USA | AY997069, NR119545 |
| *Paraglomus occultum* | IA702 ref. culture | LSU | USA | DQ273827, NG027567 |
| *Paraglomus occultum* env. sample | Not available | SSU | Not available | GCA910592205 |
| *Paraglomus pernambucanum* | Ex-type | LSU | Brazil | JX122771, JX122772 |
| *Paraglomus peruvianum* | Ex-type | SSU+ITS+LSU | Peru | MW794283-MW794291 |
| *Paraglomus peruvianum* env. sample | Not available | SSU+ITS+LSU | China | KM207870, KM207873, KM207883, KM207884, KM207885, KM207889, KM207891, KM207892, KM207905, KM207906 |
| *Paraglomus peruvianum* env. sample | Not available | SSU+ITS+LSU | Not available | EUK1269204 |
| *Paraglomus* sp.1 env. sample | Not available | SSU+ITS+LSU | South Africa | KY242680 |
| *Paraglomus* sp.2 env. Sample | Not available | SSU+ITS+LSU | China | KM207919, KM207920, KM207926, KM207932, KM207947, KM207952, KM207954, KM207956, KM207960, KM208034 |
| *Paraglomus* sp.3 | ASV281 | SSU+ITS+LSU | USA | MT765575 |
| *Paraglomus* sp.4 | ASV329 | SSU+ITS+LSU | USA | MT765623 |
| *Paraglomus* sp.5 env. sample | BG3_95 | SSU+ITS+LSU | China | KY114686 |
| *Paraglomus* sp.6 | ASV259 | SSU+ITS+LSU | USA | MT765553 |
| *Paraglomus* sp.7 | ASV219 | SSU+ITS+LSU | USA | MT765513 |
| *Paraglomus* sp.7 | ASV356 | SSU+ITS+LSU | USA | MT765650 |
| *Paraglomus* sp.8 env. sample | G4741 | SSU+ITS+LSU | Estonia | EUK1223177 |
| *Paraglomus* sp.8 env. sample | Not available | SSU+ITS+LSU | Peru | HF970206 |
| *Paraglomus* sp.9 env. sample | G5116 | SSU+ITS+LSU | Brazil | EUK1698282 |
| *Paraglomus* sp.10 env. sample | G6005 | SSU+ITS+LSU | Panama | EUK1698737 |
| *Paraglomus* sp.10 env. sample | G5676 | SSU+ITS+LSU | Kenya | EUK1698764 |
| *Paraglomus* sp.10 env. sample | S932 | SSU+ITS+LSU | Oman | EUK1698266, EUK1698552 |
| *Paraglomus* sp.10 env. sample | S824 | SSU+ITS+LSU | Tanzania | EUK1698226 |
| *Paraglomus* sp.11 env. sample | G6006 | SSU+ITS+LSU | Panama | EUK1698801 |
| *Paraglomus* sp.11 env. sample | MX24 | SSU+ITS+LSU | Mexico | EUK1698196 |
| *Paraglomus* sp.11 env. sample | G3059 | SSU+ITS+LSU | India | EUK1699013, EUK1699061 |
| *Paraglomus* sp.12 | 5088_2360A | SSU+ITS+LSU | Canada | PX215150, PX215153, PX215157 |
| *Paraglomus* sp.12 | 4423_1159D | SSU+ITS+LSU | Canada | PX215152, PX215154, PX215155, PX215156, PX215159, PX215160, PX215161 |

| | | | | |
|---|---|---|---|---|
| *Pervetustus simplex* | Ex-type | SSU+ITS+LSU | Tunisia | KY630238-KY630243, KY630245, KY630247, KY630249 |
| Environmental sample | Not available | SSU+ITS+LSU | Not available | UDB014734 |
| Environmental sample | S047 | SSU+ITS+LSU | Puerto Rico | EUK1188395, EUK1188396 |
| Environmental sample | G5040 | SSU+ITS+LSU | British Virgin Islands | EUK1188392, EUK1188393 |
| Environmental sample | S045 | SSU+ITS+LSU | Puerto Rico | EUK1231013, EUK1231018 |
| Environmental sample | Not available | SSU+ITS+LSU | Puerto Rico | EUK1104647 |
