## Supplementary material for "Revision of Paraglomeromycetes unveils a new order (Pervertustales), a new family (Innosporaceae), and a new genus (*Opikia*) in Glomeromycota": Table S1

Table S1. EUKARYOME accession numbers of environmental sequences, their origin, and percent identity to sequences of *Opikia catenata* that clustered in the clade with this new species

| Accession | Isolation source | Sampling date | Coutry/locality | Biome | Percent identity | References |
|---|---|---|---|---|---|---|
| EUK1699452 | soil | 2017-05-07 | USA, Michigan, Niagara Falls; 43.0685-79.0697 | Temperate broadleaf forest with Tsuga canadensis, Quercus rubra Prunus sp. | 97.55 | Salestani et al. unpubl. |
| EUK1699445 | soil | 2015-01-21 | Mexico, Michoacán, Cuitzeo; 19.6212 -101.3333 | Tropical broadleaf forest with Quercus castanea, Q. deserticola | 97.23 | Salestani et al. unpubl. |
| EUK1699496 | soil | 2012-12-14 | Iran, Shirgah; 36.3040 52.9010 | Temperate broadleaf forest biome with Parrotia | 96.60 | Salestani et al. unpubl. |
| EUK1699461 | soil | 2015-05-18 | USA, VA, Shenandoah, 38.8756-78.2089 599 | Temperate broadleaf forest with Quercus with Tilia, Carya, Acer, Pinus, Juglans, Ulmus | 97.42 | Salestani et al. unpubl. |
| EUK1699423 | soil | 2016-03-05 | South Africa, Marloth Nature Reserve, 1-33.9932 20.4591 | subtropical broadleaf forest | 96.79 | Salestani et al. unpubl. |
| EUK1699469 | soil | 2015-01-21 | Mexico, Michoacán, Cuitzeo, 19.6212-101.3333 2550 | Tropical broadleaf forest with Quercus castanea, Q. deserticola | 96.72 | Salestani et al. unpubl. |
| EUK1699455 | soil | 2012-12-14 | Iran, Shirgah, 36.3040 52.9010 | Temperate broadleaf forest with Parrotia | 96.16 | Salestani et al. unpubl. |
| EUK1699566 | soil | 2010-05-23 | Russia, Primorye Krai, Kedrovaya Pad W, 43.1281 131.4112 | Temperate broadleaf forest with Tilia, Quercus, Populus, Corylus, Carpinus, Betula | 96.91 | Salestani et al. unpubl. |
| EUK1699353 | soil | 2017-05-07 | USA, Michigan, Niagara Falls, 43.0685-79.0697 | Temperate broadleaf forest with Tsuga canadensis, Quercus rubra, Prunus | 97.85 | Salestani et al. unpubl. |
| EUK1631673 | soil | 2018-09-30 | Estonia, Őssu, Karja, 1058.3633 26.6557 | Village biome with Vitis vinifera, Malus domestica, Pyrus communis;Picea abies, Corylus avellana, Prunus, Ribes, Populus tremula | 96.80 | Tedersoo et al. 2024 |
